# The T-type voltage-gated Ca^2+^ channel Ca_V_3.1 as a candidate receptor for *Pasteurella multocida* toxin and contributes to the disruption of respiratory epithelial barrier induced by the toxin

**DOI:** 10.1101/2024.09.06.611746

**Authors:** Haixin Bi, Fei Wang, Lin Lin, Dajun Zhang, Menghan Chen, Yuyao Shang, Lin Hua, Huanchun Chen, Bin Wu, Zhong Peng

## Abstract

*Pasteurella multocida* toxin (PMT) is an exotoxin produced by several members of the zoonotic respiratory pathogen *P. multocida*. The role of PMT in disrupting the mammalian respiratory barrier remains to be elucidated. In this study, we discovered that inoculation of recombinantly expressed PMT increased the permeability of the respiratory epithelial barrier in mouse and respiratory cell models. This was evidenced by a decreased expression of tight junctions (ZO-1, occludin) and adherens junctions (β-catenin, E-cadherin), as well as enhanced cytoskeletal rearrangement. In mechanism, we demonstrated that PMT inoculation induced cytoplasmic Ca^2+^ inflow, leading to an imbalance of cellular Ca^2+^ homeostasis and endoplasmic reticulum stress. This process further stimulated the RhoA/ROCK signaling, promoting cytoskeletal rearrangement and reducing the expression of tight junctions and adherens junctions. Notably, the T-type voltage-gated Ca^2+^ channel Ca_V_3.1 was found to participate in PMT-induced cytoplasmic Ca^2+^ inflow. Knocking out Ca_V_3.1 significantly reduced the cytotoxicity induced by PMT on swine respiratory epithelial cells and mitigated cytoplasmic Ca^2+^ inflow stimulated by PMT. Further analysis identified Ser (aa92), Glu (aa155), Tyr (aa167), and Leu (aa448) as crucial sites utilized by PMT to interact with Ca_V_3.1. These findings suggest Ca_V_3.1 serves as an important host receptor of PMT and contributes to PMT-induced respiratory epithelial barrier disruption.

**Importance:** PMT is a significant toxin produced by the zoonotic respiratory pathogen *P. multocida*, yet little is known about its pathogenesis beyond causing progressive atrophic rhinitis in pigs. In our study, we have discovered that PMT has the capacity to disrupt the integrity of the mammalian respiratory epithelial barrier. This disruption involves an imbalance in cellular Ca^2+^ homeostasis, endoplasmic reticulum stress, and activation of the RhoA/ROCK signaling pathway induced by PMT. Importantly, we have identified Ca_V_3.1 as a pivotal receptor that plays a crucial role in the pathogenic effects of PMT. Our findings highlight the potential of Ca_V_3.1 as a target for intervention strategies aimed at combating the detrimental effects of PMT.

## Introduction

*Pasteurella multocida* strains are zoonotic and can infect a wide range of mammals, including pigs, dogs, rabbits, rodents, and humans (1). In agriculture, infections caused by *P. multocida* has resulted in significant economic losses in farm animals. For example, *P. multocida* can induce progressive atrophic rhinitis (PAR), a chronic wasting disease observed in pigs globally (2). This disease is estimated to cause a 5% decrease in growth during fattening, adding approximately $1.39 in production costs per slaughtered pig (3). *P. multocida* strains associated with PAR commonly produce a 146 kDa exotoxin known as *Pasteurella multocida* toxin (PMT) (4). This toxin consists of two domains: the N-terminus and the C-terminus. The N-terminus is primarily responsible for binding to host cell receptors, facilitating the translocation of the toxic active C-terminal domain into the cytoplasm, releasing the toxic region of the C-terminus, and thereby activating relevant signaling pathways to exert its toxic effects (5, 6). Early studies indicate that PMT uses gangliosides, such as GM1, GM2, and GM3, as receptors (7). More recently, the LDL Receptor Related Protein1 (LRP1) has also been identified as a key receptor for PMT through CRISPR-Cas9-based genome-wide screening (8). However, there might be additional host factors on the cell surface that contribute to PMT exerting its effects, and further exploration of the interactions between PMT and these host factors is needed.

The respiratory tract constitutes the primary line of defense against respiratory pathogen infections, encompassing physical, immune, and chemical barriers crucial for upholding host homeostasis (9). The fundamental structure of the respiratory epithelial barrier comprises epithelial cells interconnected by tight junctions and adherens junctions between adjacent cells (10). Previous studies have suggested that variations in the quantity and continuity of tight junctions and adherens junctions influence the permeability of the epithelial barrier (11, 12). Disruption of barrier function arises when the cell cytoskeleton undergoes remodeling, resulting in the altered localization of ZO-1, a critical tight junction protein (13). RhoA facilitates the bundling of filamentous actin (F-actin) into stress fibers, leading to the redistribution of tight junction proteins and barrier disruption (14). ROCK, a downstream effector of RhoA, plays a role in cytoskeletal adjustment and cell contraction (15). While it is established that PMT can activate the RhoA/ROCK signaling pathway (16, 17), there is limited understanding of whether PMT can induce dysfunction in the respiratory epithelial barrier and the underlying mechanisms.

In this study, we explore the impact of PMT on the host respiratory epithelial barrier using mouse and cell models. Our findings reveal that PMT also disrupts the mammalian respiratory epithelial barrier, with this process involving the activation of RhoA/ROCK signaling induced by the PMT-stimulated imbalance in cellular Ca^2+^ homeostasis. Remarkably, we demonstrate that the T-type voltage-gated Ca^2+^ channel Ca_V_3.1, known as a receptor for *Bordetella* dermonecrotic toxin (18), could interact with PMT and play a role in this process.

## Results

### PMT inoculation increases respiratory epithelial barrier in mouse models

To investigate the impact of PMT on the mammalian respiratory epithelial barrier, experimental mice were intranasally inoculated with recombinant PMT (30 μg per mouse) or PBS (Fig. 1A, B). At 48 hours post-inoculation (hpi), vascular permeability in the trachea and lungs was evaluated by intravenously injecting Evans blue dye via the tail vein, following established protocols (11). The experimental findings revealed a notable increase in vascular permeability in the trachea and lungs of mice exposed to PMT compared to the control group treated with PBS (Fig. 1C). Histopathological and transmission electron microscopy examinations unveiled significant tissue damage in the tracheas and lungs of mice from the PMT-treated group (Fig. 1D, E). Utilizing a monoclonal antibody specific to PMT for the detection of PMT in lung tissues of mice treated with PMT/PBS, PMT protein was identified in the lungs of mice in the PMT-treated group (Fig. 1F). Western blot analysis assessing the expression levels of the crucial tight junction protein ZO-1 exhibited a marked decrease in expression levels in the PMT-treated group of mice compared to the PBS group (Fig. 1G). These outcomes suggest that PMT may induce the disruption of the respiratory epithelial barrier by downregulating the expression of tight junction proteins.

**Fig. 1.**
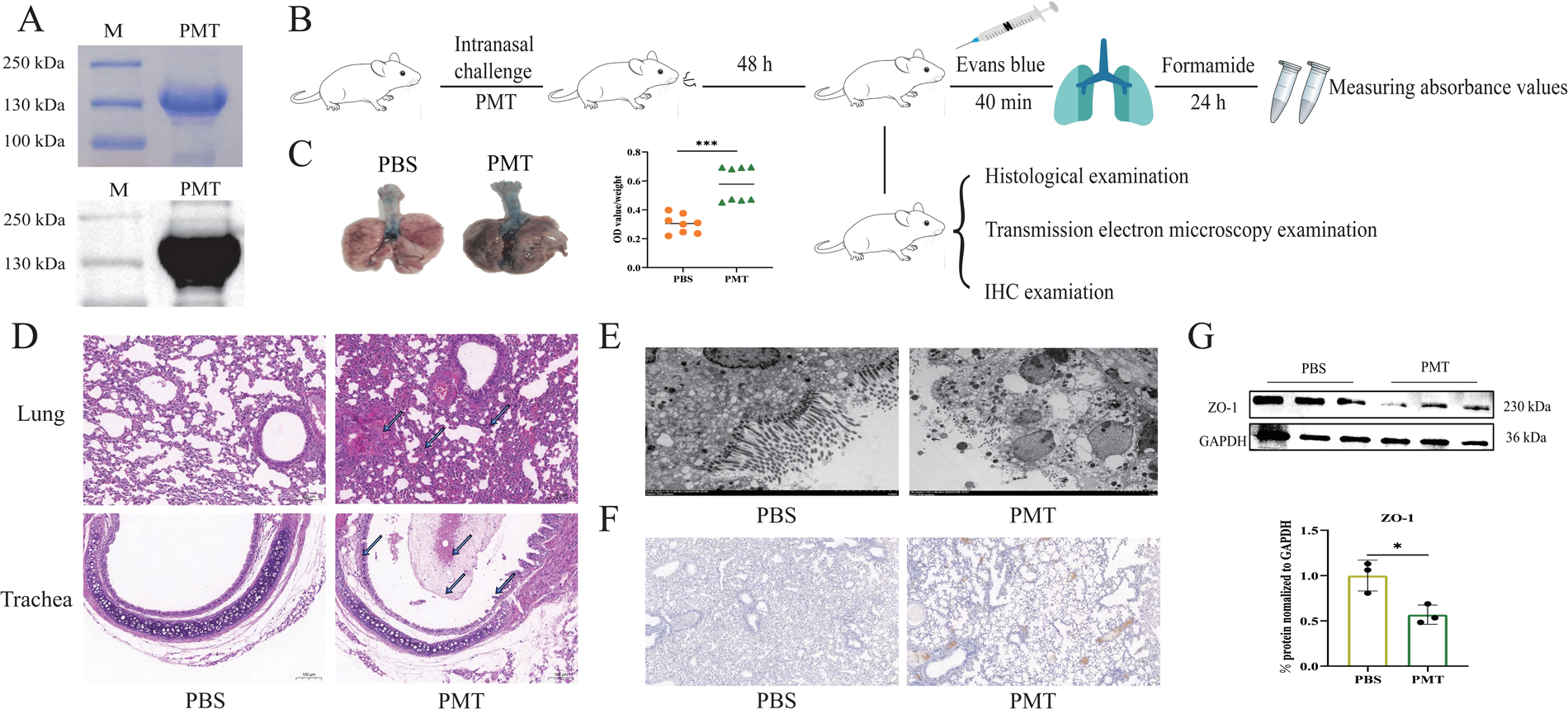
*In vivo* tests conducted in mouse models to evaluate the impact of *Pasteurella multocida* toxin (PMT) on the mammalian respiratory epithelial barrier. **(A)** SDS-PAGE (blue band) and western blot (black band) analyses of recombinantly expressed PMT protein using the *Escherichia coli* expression system. **(B)** Study design of the mouse experiments. Experimental mice were intranasally inoculated with PMT/PBS. To assess the permeability of the lungs and trachea, each mouse was intravenously injected with Evans blue dye 48 hours post-treatment. After 40 minutes, the mice were euthanized, and the dye was extracted from the lungs and trachea using formamide, followed by measuring the absorbance at OD_620_. Additionally, murine lungs and trachea were also examined through immunohistochemistry, histological staining, and electron microscopy observations. **(C)** Visualizations of the tracheae and lungs obtained from PMT and PBS treated mice, along with the quantification of Evans blue in the tracheae and lungs. **(D)** Histological examinations of tracheae and lungs obtained from PMT and PBS treated mice (scale bars = 100 μm). Lung damages in PMT-treated mice include thickening of alveolar walls, neutrophil infiltration, and thrombosis. Tracheal damages encompass epithelial cell shedding, bleeding, cytoplasmic vacuolation, and pale pink foam-like secretion within the lumen. **(E)** Ultrastructural damages in the trachea of PMT-inoculated mice observed via electron microscopy, characterized by disorganized epithelial cell arrangement, severe vacuolization of cells, and enlarged intercellular gaps. **(F)** Immunohistochemistry analysis demonstrates the presence of PMT in the lungs of PMT-treated mice. **(G)** Western blot analysis indicated a decrease in ZO-1 expression in the lungs of PMT-inoculated mice compared to PBS-treated mice. The data are represented as mean ± SD, with the significance level set at *P* < 0.05 (∗).

### PMT inoculation decreases tight junctions and adherens junctions between neighboring respiratory epithelial cells

We next conducted dextran-based trans-well permeability assays to explore the effect of PMT on the respiratory epithelial barrier (Fig. 2A). To ensure the validity of subsequent experiments, we initially assessed cell viability before proceeding with *in vitro* studies. The results demonstrated that the treatment of MLE-12 (murine respiratory epithelial cells) or NPTr (swine respiratory epithelial cells) cells with 25 µg/mL of PMT for 48 hours did not significantly impact cell viability (Fig. 2B). The dextran-based transwell permeability assays showed that treating MLE-12/NPTr cells with 25 µg/mL of PMT resulted in a decline in the integrity of the cell barrier function (Fig. 2C). A reduction in the expression of tight junctions (ZO-1 and occludin) and adherens junctions (E-cadherin, β-catenin) was observed in PMT-treated respiratory epithelial cells compared to cells treated with PBS (Fig. 2D, E, F). These observations align with the results from the mouse experiments, suggesting that PMT disrupts the respiratory epithelial barrier by downregulating the expression of tight junction proteins and adherens junctions.

**Fig. 2.**
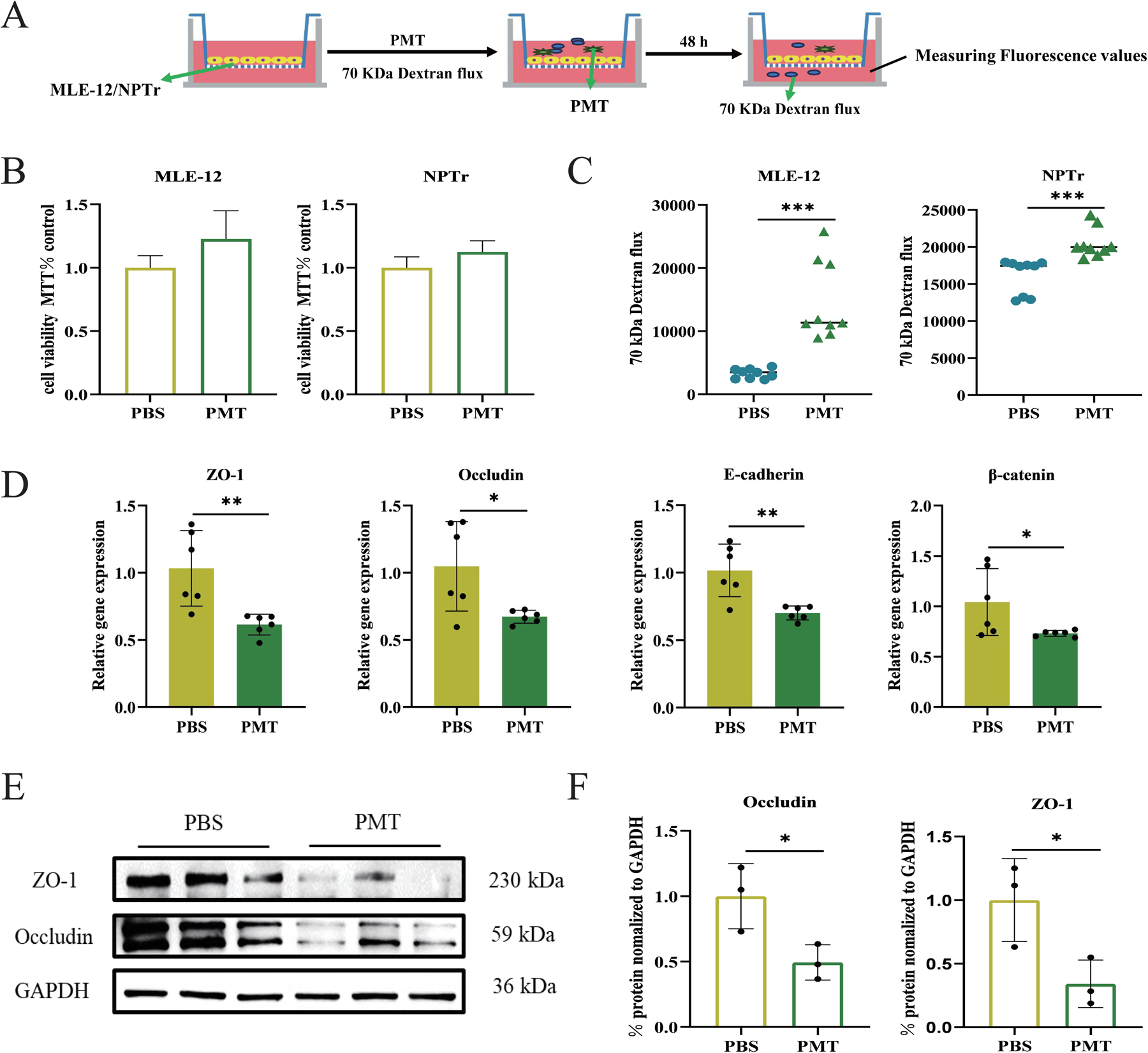
*In vitro* tests conducted in mammalian respiratory epithelial cell models to evaluate the impact of *Pasteurella multocida* toxin (PMT) on the barrier functions formed by the swine (NPTr) and/or murine (MLE-12) respiratory epithelial cells. **(A)** Study design of the dextran-based trans-well permeability assays. **(B)** Cytotoxicity assays revealing the conditions of NPTr/MLE-12 at 48 h post PMT inoculation. **(C)** Dextran-based trans-well permeability assays showing the increase in the vascular permeability of NPTr and/or MLE-12 cells induced by PMT. **(D)** The transcription levels of ZO-1, occludin, E-cadherin, and β-catenin in PMT-treated cells compared to PBS-treated cells. **(E)** Expression of ZO-1 and occludin in PMT-treated cells compared to PBS-treated cells. **(F)** Quantifications of the western blot results (panel E) from cell extracts in triplicate. Data represent mean ± SD. The significance level was set at *P* < 0.05 (∗), *P* < 0.01 (∗∗), or *P* < 0.001 (∗∗∗).

### PMT induces respiratory epithelial barrier disruption by activating the RhoA/ROCK signaling pathway, leading to cellular cytoskeletal rearrangements

In the pursuit of further understanding the mechanism by which PMT disrupts the respiratory epithelial barrier, immunofluorescence experiments were conducted. In cells treated with PBS, ZO-1 was predominantly situated at the cell periphery, displaying a continuous distribution. F-actin exhibited a circular and continuous distribution at the edges of NPTr cells, devoid of visible stress fibers (Fig. 3A). Upon PMT treatment, the expression of ZO-1 decreased, transitioning from a continuous to a discontinuous distribution state. The quantity of F-actin at the cell periphery declined, while stress fibers increased and were notably observed in the cell center (Fig. 3A). Analysis of RhoA in mice and NPTr cells treated with PMT/PBS revealed a substantial increase in RhoA expression in the lung tissues of mice post PMT treatment (Fig. 3B). The expression levels of RhoA in NPTr cells escalated in a time-dependent manner, with the most pronounced increase in RhoA and ROCK1 protein expression observed after 48 hours of treatment (Fig. 3C).

**Fig. 3.**
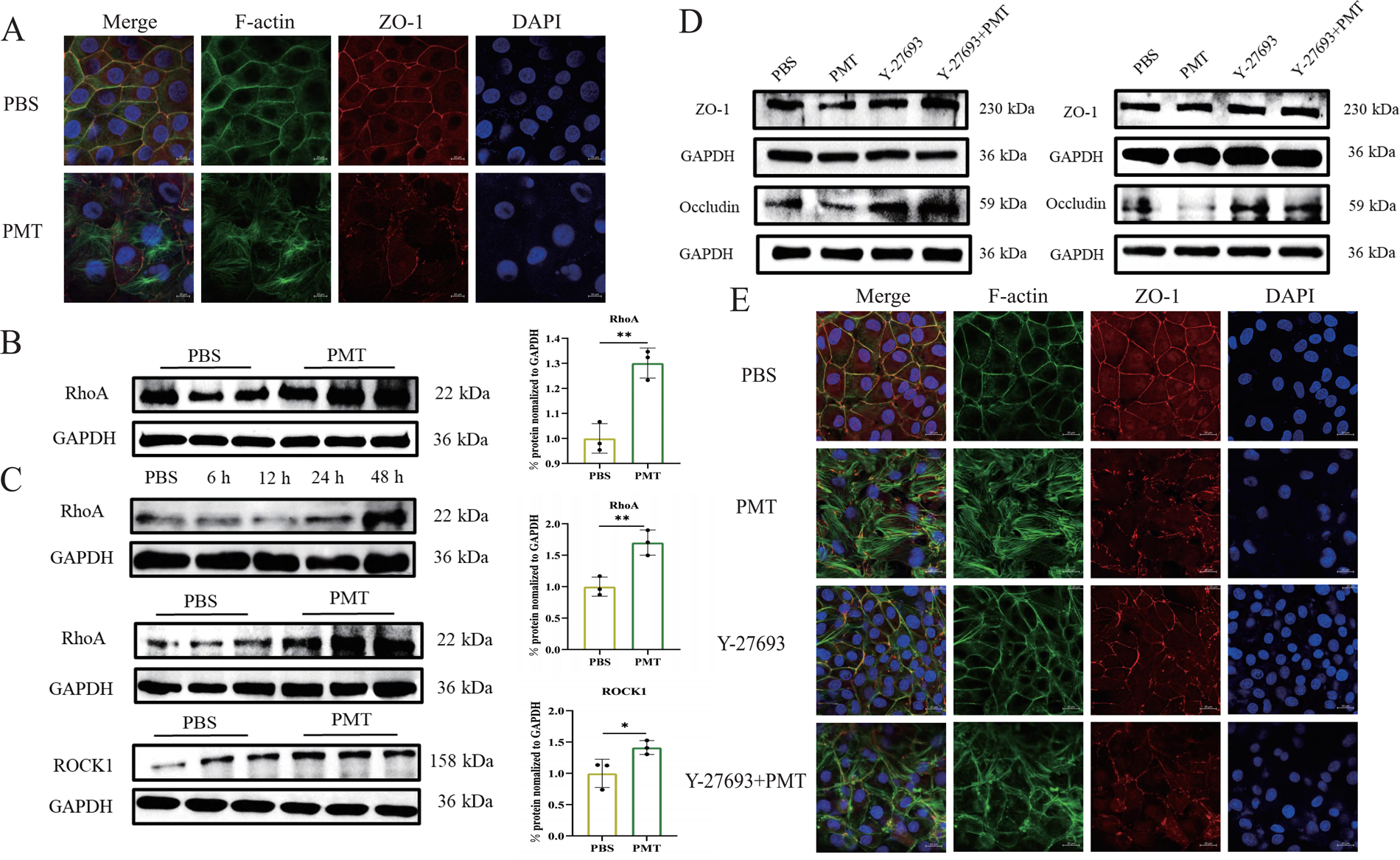
*Pasteurella multocida* toxin (PMT) disrupts the respiratory epithelial barrier by activating the RhoA/ROCK signaling pathway, leading to cytoskeletal rearrangement. **(A)** Immunofluorescence images showing reduced and disorganized F-actin and ZO-1 in PMT-treated NPTr cells compared to the PBS-treated cells. Scale bar: 20 µm. **(B)** The expression of RhoA in the lungs of mice inoculated with PMT compared to the PBS-treated mice. **(C)** The expression of RhoA and ROCK1 in NPTr cells inoculated with PMT at different time points compared to the PBS-treated cells. **(D)** The expression of ZO-1 and occludin in NPTr cells treated with PMT, PMT and the ROCK-specific inhibitor Y-27632, the ROCK-specific inhibitor Y-27632, and PBS. **(E)** Immunofluorescence images showing reduced and disorganized F-actin and ZO-1 in NPTr cells treated with PMT, PMT and the ROCK-specific inhibitor (Y-27632), the ROCK-specific inhibitor (Y-27632), and PBS. Data represent mean ± SD. The significance level was set at *P* < 0.05 (∗), or *P* < 0.01 (∗∗).

To delve into the role of the RhoA/ROCK signaling pathway in the PMT-induced disruption of the respiratory epithelial barrier, MLE-12/NPTr cells were pre-treated with the ROCK inhibitor Y-27693. Western blot outcomes indicated that following pretreatment with Y-27693, the protein expression levels of ZO-1 and occludin in the inhibitor-treated group were significantly higher than those in the group without inhibitor treatment (Fig. 3D). Immunofluorescence results demonstrated that after pretreatment with the ROCK-specific inhibitor Y-27693, ZO-1 partially reinstated its continuous distribution state, with a notable increase in F-actin distributed at the cell periphery while stress fiber formation markedly decreased (Fig. 3E).

### PMT induces imbalance of cellular Ca^2+^ homeostasis, leading to endoplasmic reticulum stress and RhoA activation

Given that calcium ions (Ca^2+^) have been reported to play a role in regulating the RhoA/ROCK signaling pathway (13), we hypothesized that Ca^2+^ might be involved in the upregulation of RhoA stimulated by PMT. To investigate this, we labeled Ca^2+^ in NPTr cells using a calcium ion fluorescent probe Fluo-3 AM and examined the changes in Ca^2+^ levels following PMT treatment. The results revealed a significant increase in the fluorescence intensity of cytoplasmic Ca^2+^ after 6 to 48 hours of treatment with 25 µg/mL of PMT, suggesting that PMT could induce cellular Ca^2+^ homeostasis imbalance (Fig. 4A). Previous studies have also highlighted a relationship between cytoplasmic Ca^2+^ influx and endoplasmic reticulum stress (19, 20). Therefore, we further investigated the changes in endoplasmic reticulum stress-related markers after PMT treatment of NPTr cells. At the gene level, the expression levels of endoplasmic reticulum stress-related marker genes showed an increasing trend after PMT stimulation for 6, 12, and 48 hours, but endoplasmic reticulum stress was suppressed at 24 hours (Fig. 4B). At the protein level, the expression levels of GRP78, IRE1, and p-IRE1 increased after treatment with 25 µg/mL of PMT for 6 to 48 hours (Fig. 4C). Through quantitative real-time PCR (qPCR) assays, we also found that the gene expression levels of GRP78, PERK, and IRE1 decreased after using the calcium chelator BAPTA-AM (Fig. 4D). To investigate the role of endoplasmic reticulum stress in RhoA activation, cells were pretreated with the endoplasmic reticulum stress inhibitor 4-PBA. Western blot showed that the RhoA protein expression level in the 4-PBA treated group was significantly downregulated compared to the control cells post PMT treatment (Fig. 4E). The above findings indicate that PMT can lead to cytoplasmic Ca^2+^ influx and endoplasmic reticulum stress, and endoplasmic reticulum stress plays a role in the process of PMT-induced elevation of RhoA.

**Fig. 4.**
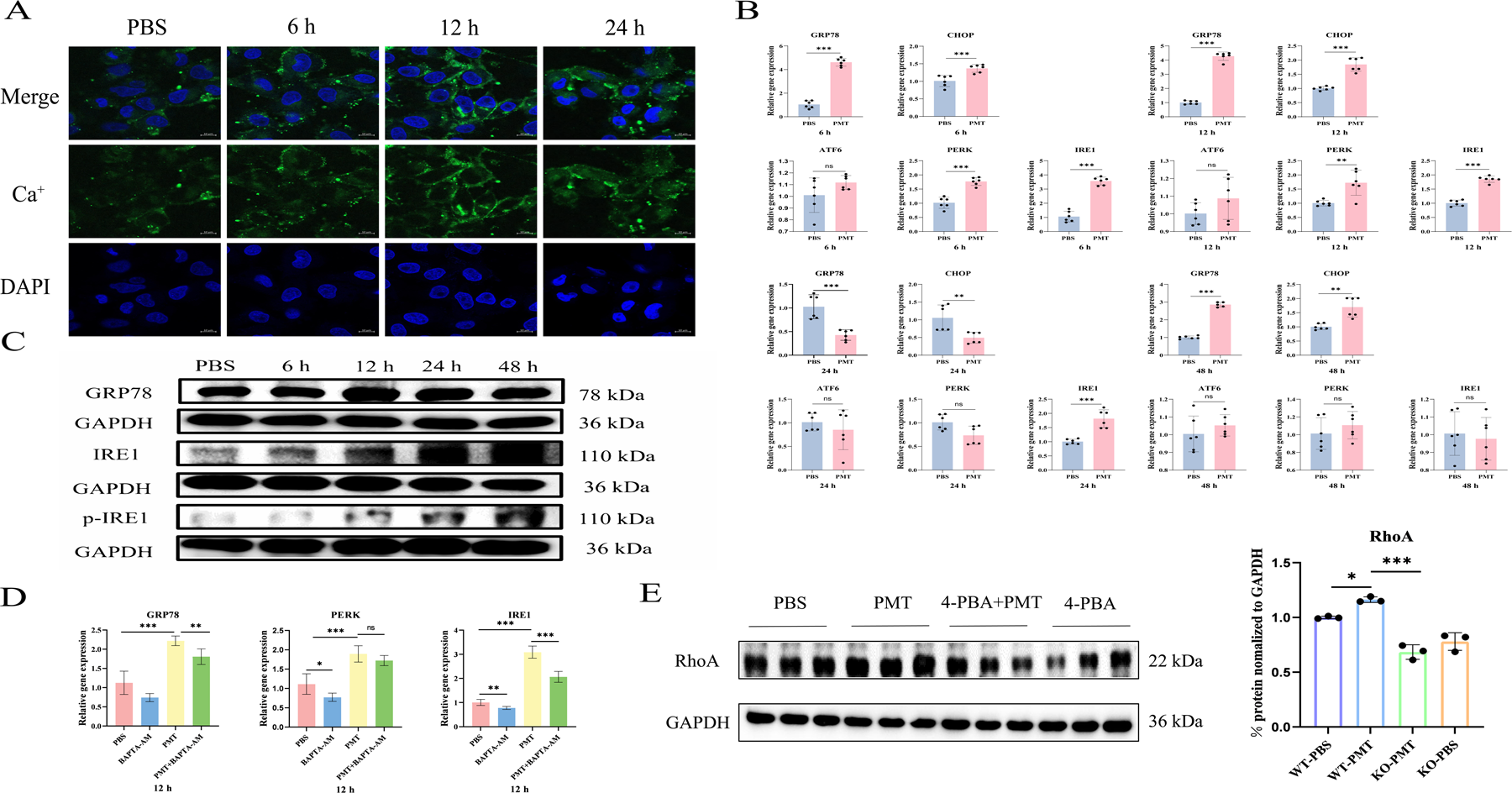
*Pasteurella multocida* toxin (PMT) inoculation stimulates cellular calcium (Ca^2+^) homeostasis imbalance and endoplasmic reticulum stress. **(A)** Immunofluorescence images showing cellular Ca^2+^ (labeled with the calcium ion fluorescent probe Fluo-3 AM, green) in PMT-treated NPTr cells at different time points compared to the PBS-treated cells. Scale bar: 10 µm. **(B)** The transcription levels of different endoplasmic reticulum stress markers (GRP78, CHOP, ATF6, PERK, IRE1) in PMT-treated NPTr cells at different time points compared to the PBS-treated cells. **(C)** The expressions of different endoplasmic reticulum stress markers (GRP78, IRE1, and phosphorylated IRE1[p-IRE1]) in PMT-treated NPTr cells at different time points compared to the PBS-treated cells. **(D)** The transcription levels of different endoplasmic reticulum stress markers (GRP78, PERK, IRE1) in NPTr cells treated with PMT, PMT and the Ca^+^ chelating agent (BAPTA-AM), the Ca^+^ chelating agent (BAPTA-AM), and PBS for 12 h. **(E)** The expression of RhoA in NPTr cells treated with PMT, PMT and the endoplasmic reticulum stress inhibitor (4-PBA), the endoplasmic reticulum stress inhibitor (4-PBA), and PBS. Data represent mean ± SD. The significance level was set at *P* < 0.05 (∗), *P* < 0.01 (∗∗), or *P* < 0.001 (∗∗∗).

### The T-type voltage-gated Ca^2+^ channel Ca_V_3.1 could interact with PMT and involves in PMT-induced cellular calcium homeostasis imbalance and respiratory barrier dysfunction

A recent study has identified the T-type voltage-gated Ca^2+^ channel Ca_V_3.1 (encoded by *Cacna1g*) as a candidate receptor through which *Bordetella* dermonecrotic toxin (bDNT) (18). Since PMT shares similarities with bDNT at both N- and C-terminals (6), we next investigate whether Ca_V_3.1 is involved in PMT-induced cellular calcium imbalance. To achieve this, we generated a Ca_V_3.1-KO NPTr cells using CRISPR-Cas9 system. Cytotoxicity tests revealed that Ca_V_3.1-KO significantly decreased reduced PMT-induced cytotoxicity (Fig.S1 in supplementary materials). Subsequent analysis of cytoplasmic Ca^2+^ levels post PMT treatment revealed a significant inhibition of Ca^2+^ influx in Ca_V_3.1-KO cells compared to wild-type cells (Fig. 5A). qPCR and western blot results indicated a suppression of endoplasmic reticulum stress in Ca_V_3.1-KO cells (Fig. 5B, C). Moreover, RhoA protein levels were notably lower in Ca_V_3.1-KO cells, and immunofluorescence results showed decreased stress fibers and partial ZO-1 recovery in these cells following PMT stimulation (Fig. 5D, E). These findings collectively suggest that Ca_V_3.1 plays a crucial role in PMT’s toxic effects.

**Fig. 5.**
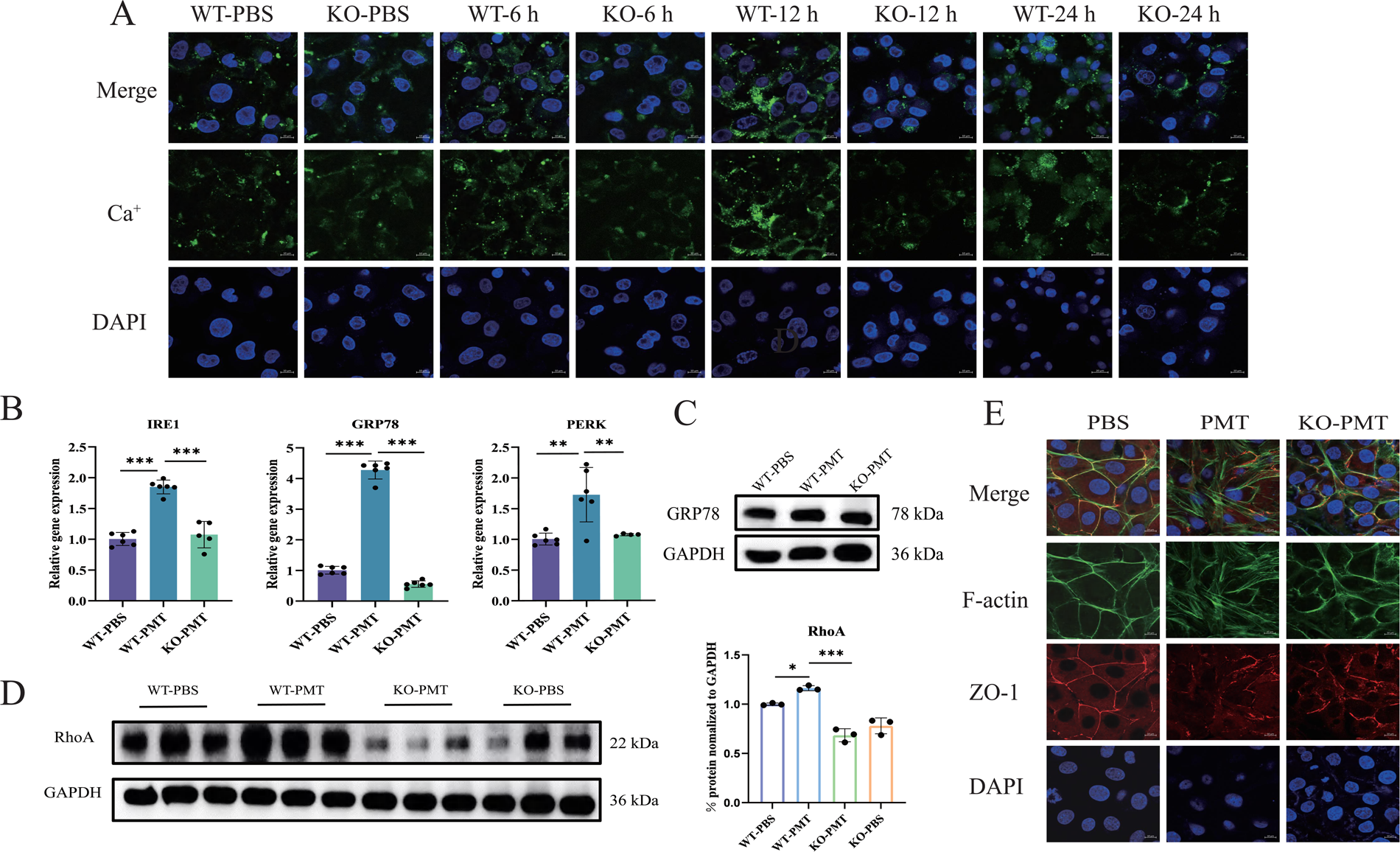
Impacts of Ca_v_3.1 on *Pasteurella multocida* toxin (PMT) induced respiratory epithelial barrier disruption. **(A)** Immunofluorescence images showing cellular Ca^2+^ (labeled with the calcium ion fluorescent probe Fluo-3 AM, green) in NPTr wild type and Ca_v_3.1-KO cells treated by PMT at different time points compared to the PBS-treated cells. **(B)** The transcription levels of different endoplasmic reticulum stress markers (GRP78, PERK, IRE1) in PMT-treated Ca_v_3.1-KO cells compared to PMT-treated wild type cells and PBS-treated cells. **(C)** The expression of endoplasmic reticulum stress marker GRP78 in PMT-treated Ca_v_3.1-KO cells compared to PMT-treated wild type cells and PBS-treated cells. **(D)** The expression of RhoA in PMT-treated Ca_v_3.1-KO cells compared to PMT-treated wild type cells and PBS-treated cells. **(E)** Immunofluorescence images showing reduced and disorganized F-actin and ZO-1 in PMT-treated Ca_v_3.1-KO cells compared to PMT-treated wild type cells and PBS-treated cells. Data represent mean ± SD. The significance level was set at *P* < 0.05 (∗), *P* < 0.01 (∗∗), or *P* < 0.001 (∗∗∗).

To explore the interaction between PMT and Ca_V_3.1, structural models of PMT were generated using AlphaFold3 (21), and molecular docking analysis revealed that PMT can dock with Ca_V_3.1 (protein data bank [PDB] ID: O43497 (22)) (Fig. 6A). The calculated binding energy was -84.46 kcal/mol, with key docking amino acids in PMT identified as Ser (aa92), Glu (aa155), Tyr (aa167), and Leu (aa448). A co-immunoprecipitation (co-IP) assay confirmed the binding between PMT and CaV3.1 (Fig. 6B). Subsequent mutations targeting specific amino acid residues in PMT, including Tyr (aa100), Asp (aa149), His (aa150), and Tyr (aa152), significantly decreased the cytotoxicity induced by PMT, suggesting the importance of these residues in PMT-induced cytotoxicity in respiratory epithelial cells (Fig. 6C, D).

**Fig. 6.**
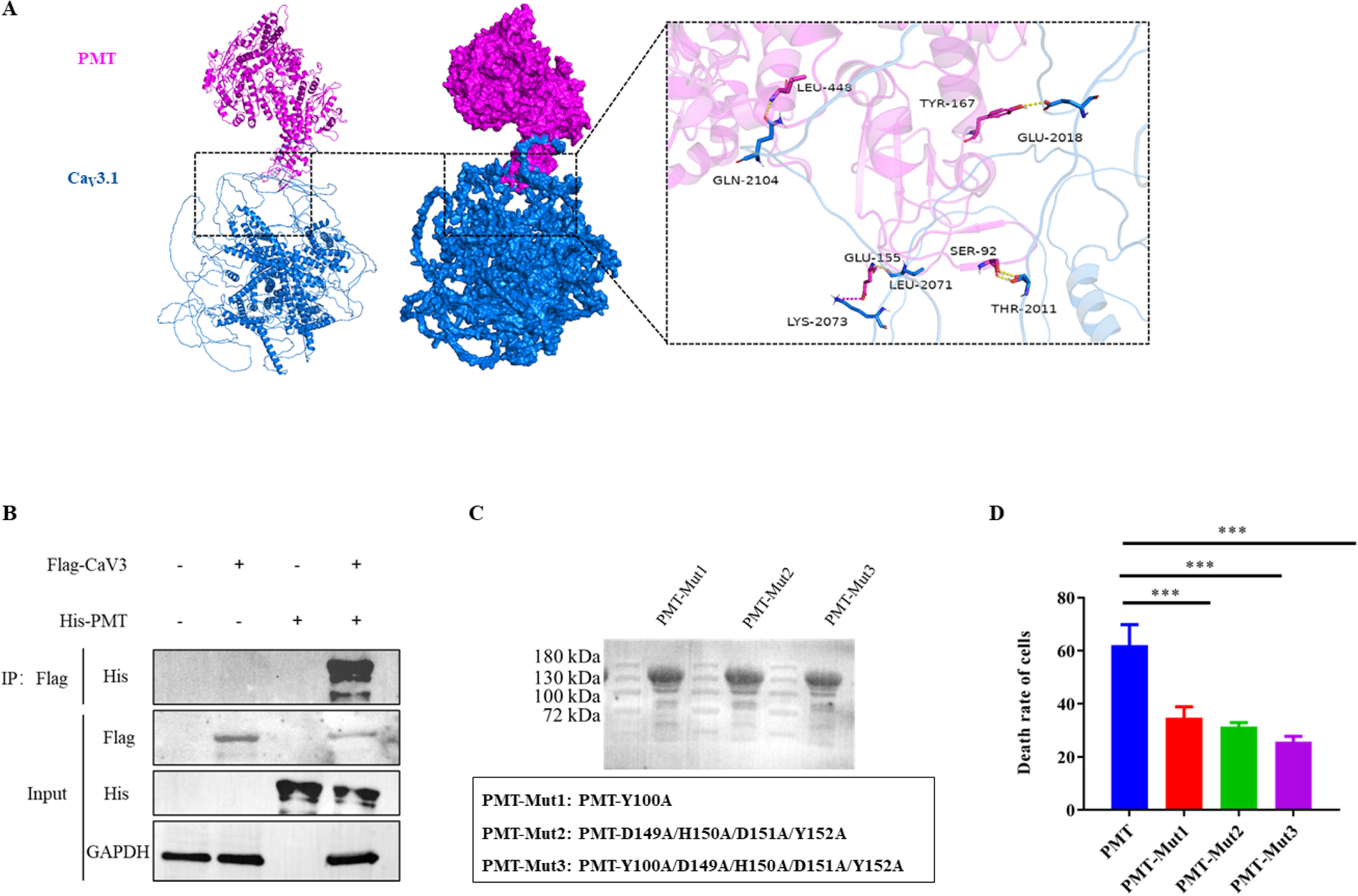
Interactions between *Pasteurella multocida* toxin (PMT) and Ca_v_3.1. **(A)** Molecular docking analysis demonstrates the interaction between PMT and Ca_v_3.1. **(B)** Co-immunoprecipitation verifies the interaction between PMT and Ca_v_3.1. **(C)** The expression of different proteins generated from PMT is shown, where mutations of amino acids at various positions in the toxin were performed. Amino acids selected for mutation are indicated within a black frame. **(D)** Susceptibility of NPTr to PMT and generated proteins. Data represent mean ± SD. The significance level was set at *P* < 0.001 (∗∗∗).

## Discussion

As the sole exotoxin encoded by *P. multocida*, a zoonotic respiratory pathogen, PMT is known to cause PAR (4). However, its role in disrupting the mammalian respiratory barrier in remains to elucidated. In this study, we demonstrated that PMT exhibited an effect on disrupting the mammalian respiratory epithelial barrier, as evidenced by assessments in both mouse and respiratory epithelial cell models. In mouse models, intranasal inoculation of PMT led to a higher level of Evans Blue dye in the trachea and lungs. Evans Blue is a commonly used azo dye formulation with high affinity for plasma albumin in the blood, frequently used as a tracer to indicate the integrity of the physiologic barriers, including the respiratory epithelial barrier (11, 12). The increased level of Evans Blue dye observed in the trachea and lungs of mice inoculated with PMT indicates an enhanced permeability of the respiratory epithelial barrier. This impact was further confirmed by dextran-based trans-well permeability assays conducted in respiratory epithelial cells from mice and pigs, which is also a strategy frequently implemented for evaluating the disruption of barrier function formed by epithelial cells (11, 12). Additionally, the decrease in tight junctions (ZO-1, occludin) and adherens junctions (β-catenin, E-cadherin) induced by PMT also supports the role of this toxin in disrupting the respiratory epithelial barrier, as both tight junctions and adherens junctions between neighboring epithelial cells are crucial for maintaining tissue barrier integrity (23, 24).

Our analysis conducted in this study demonstrated that PMT stimulated cytoskeletal rearrangement and changes in ZO-1 redistribution in mammalian respiratory epithelial cells. These findings agree well with previous studies (25, 26). It should be noted that the F-actin cytoskeleton has been showed to play a central role in endothelial tight junction barrier regulation (27). The tight junction protein ZO-1 establishes a connection between the transmembrane protein occludin and the actin cytoskeleton, thus forming a stable tight junction complex crucial for maintaining the barrier function of epithelial or endothelial cells (28). Furthermore, our analysis also revealed that the RhoA/ROCK signaling was involved in PMT-induced cytoskeletal rearrangement and changes in ZO-1 redistribution. Previous studies have reported the participation of the RhoA/ROCK signaling in disrupting blood-brain barrier permeability and intestinal barrier permeability (29–31). It is not surprising that the RhoA/ROCK signaling was involved in PMT-induced respiratory epithelial barrier disruption, as this signaling is one of the most classic pathways stimulated by PMT (17). Consistently, inhibiting the RhoA/ROCK signaling rescued the disruption of the barrier function in mammalian respiratory epithelial cells, as observed in this study.

In this study, we identified Ca_V_3.1 as a potential receptor for PMT and partially elucidated its role in the disruption of the mammalian respiratory barrier induced by PMT. Notably, Ca_V_3.1 has been identified as a host receptor for *Bordetella* DNT in a recent study (18). Knocking out Ca_V_3.1 significantly reduced the cytotoxicity induced by *Bordetella* DNT on MC3T3-E1 cells (18). Similarly, we also observed a significant reduction in cytotoxicity induced by PMT on NPTr cells upon knocking out Ca_V_3.1. It has been noted that PMT shares similarities with *Bordetella* DNT at both the N-terminal region, where the receptor binding domain is located, and the C-terminal region, where the activity domain is situated (6, 7). As expected, we identified the N-terminal part of PMT as the binding domain to Ca_V_3.1. The key animo acids identified (Ser [aa92], Glu [aa155], Tyr [aa167], and Leu [aa448]) were all situated within the N-terminal 1-580 animo acid segment, which has been suggested as the receptor binding domain of PMT (6, 7).

As a T-type voltage-gated Ca^2+^ channel, Ca_V_3.1 plays a crucial role in facilitating Ca^2+^ influx into the cytosol in response to changes in membrane potential (22). Consistent with its fundamental biological function, our analysis demonstrated that knocking out Ca_V_3.1 significantly rescued Ca^2+^ influx stimulated by PMT and subsequent processes linked to the disruption of cellular Ca^2+^ homeostasis, including the upregulation of endoplasmic reticulum stress (32) and RhoA activation (33). This rescue was supported by decreased expressions of endoplasmic reticulum stress markers (IRE1, GRP78, PERK) and RhoA in in Ca_V_3.1-KO cells compared to the wild type cells following PMT inoculation. Remarkably, the knockout of Ca_V_3.1 notably reversed the cytoskeletal rearrangement (as indicated by F-actin) and ZO-1 reorganization triggered by the PMT-induced cellular Ca^2+^ imbalance, endoplasmic reticulum stress, and RhoA signaling axis, underscoring the significant role of CaV3.1 in the pathogenesis of PMT.

In conclusion, our study illustrates that PMT has the capacity to compromise the integrity of the mammalian respiratory epithelial barrier. This mechanism is intricately linked to the cellular Ca^2+^ imbalance, endoplasmic reticulum stress, and RhoA signaling axis provoked by PMT. Significantly, our research has pinpointed Ca_V_3.1 as a pivotal host receptor for PMT, highlighting its essential involvement in PMT-induced disruption of the mammalian respiratory barrier through the cellular Ca^2+^ imbalance, endoplasmic reticulum stress, and RhoA signaling axis (Fig. 7).

**Fig. 7.**
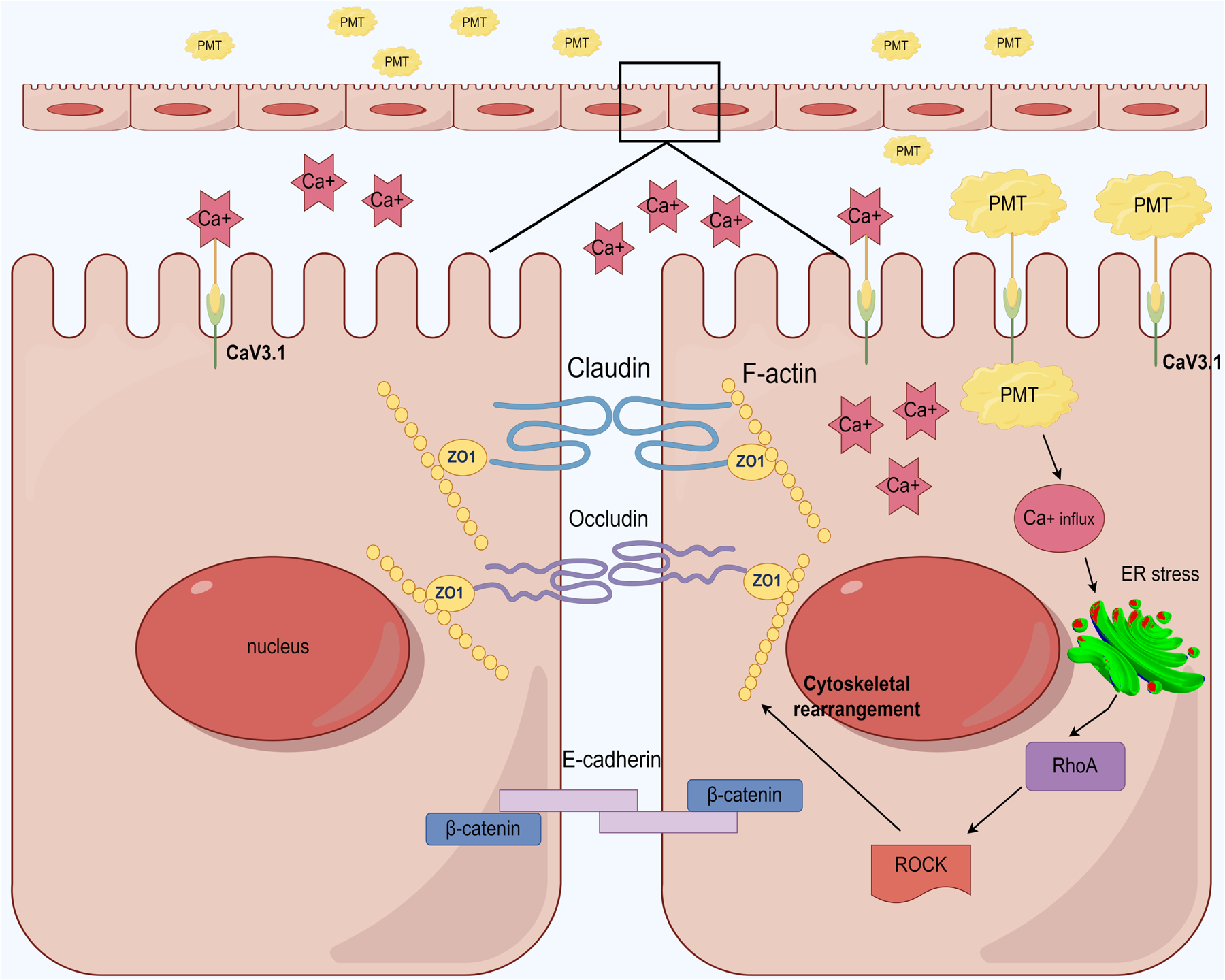
Proposed mechanisms of mammalian respiratory barrier disruption induced by *Pasteurella multocida* toxin (PMT). PMT binds to and interacts with Ca_V_3.1, promoting cytoplasmic Ca^2+^ inflow, which results in an imbalance of cellular Ca^2+^ homeostasis and endoplasmic reticulum stress. This process further stimulates RhoA/ROCK signaling, ultimately enhancing cytoskeletal rearrangement and reducing the expression of tight junctions and adherens junctions.

## Materials and methods

### Bacterial strains, cell lines, and cultural conditions

*P. multocida* HN06 (GenBank accession no. CP003313) used in this study was a capsular type D strain that produces PMT. It was isolated from a pig with PAR in Hainan province, China (34). NPTr, MLE-12, and HEK293T cells were cultured in Dulbecco’s modified eagle medium (DMEM; Gibco, Waltham, US) supplemented with 10% fetal bovine serum (FBS; Sigma-Aldrich, St. Louis, US) at 37 °C under a 5% CO_2_ atmosphere.

### Protein expression and purification

To obtain the recombinant proteins of PMT, the full-length coding gene of PMT (*toxA*) from *P. multocida* HN06 was cloned into the plasmid pET-28a using the primers listed in Table S1 in the supplementary materials. The Mut Express II Fast Mutagenesis Kit V2 (Vazyme, Nanjing, China) was used to introduce mutations in the amino acids at positions 100 (Y→A), 149 (D→A), 150 (H→A), 151 (D→A), and 152 (Y→A) in PMT, resulting in the generation of recombinant plasmids with the desired mutations. Sanger sequencing was performed to confirm the correctness of all generated plasmids. The different recombinant plasmids were then transformed into *E. coli* BL21, and the expression of different proteins was induced using 0.4 mM isopropyl-β-D-thiogalactoside (IPTG; Sigma-Aldrich, St. Louis, US). The recombinant proteins were purified using a GE Healthcare HisTrap HP nickel column (GE HealthCare, Chicago, US). Purified proteins were dialyzed at 4 °C for 24 hours using a MW14000 dialysis membrane (Biosharp, Hefei, China). Subsequently, endotoxins were removed using a commercial EtEraserTM Endotoxin Removing Kit (Bioendo, Ensenada, Mexico). The protein concentrations were determined using a commercial Enhanced BCA Protein Assay Kit (Beyotime, Shanghai, China). The recombinant proteins were stored at -80 °C for further use.

### Mouse experiments and ethic statement

Mouse experiments were conducted in Laboratory Animal Center at Huazhong Agricultural University (Wuhan, China) and were approved by the University Ethics Committee (approval ID: HZAUMO-2024-0193). Throughout the experiments, all animal handling procedures adhered to the ARRIVE guidelines 2.0 (35). Briefly, 6-week-old female specific pathogen free (SPF) Kunming (KM) mice were divided into two groups, each consisting of 6 mice. One group of mice received nasal inoculation with PMT (30 μg per mouse), while the other group was nasally administrated with PBS (50 μl per mouse). At 48 hpi, three mice were randomly selected from each group for intravenous injection of Evans blue dye (Sigma-Aldrich, St. Louis, US) via the tail vein (Sigma-Aldrich, St. Louis, US) via the tail vein. After 40 minutes, the mice were euthanized, and dye extraction from the trachea and lungs was conducted overnight using 2 mL of formamide at 55°C. Subsequently, absorbance at 620 nm was measured to evaluate vascular permeability in the murine trachea and lungs, following established protocols (11). Additionally, the remaining three mice from each group, not subjected to Evans Blue injection, were also euthanized for immunohistochemistry, histological staining, and electron microscopy observations of the lungs and trachea.

### Dextran-based *trans*-well permeability assay

A dextran-based *trans*-well permeability assay was conducted to assess the impact of PMT on the barrier function of mammalian respiratory epithelial cells, following established procedures (11, 12). Initially, approximately 1×10^5^ NPTr or MLE-12 cells were seeded onto cell culture inserts (Labselect, Hefei, China), which were positioned in the wells of a 24-well plate (Corning, USA). These cells were cultured in antibiotic-free DMEM medium supplemented with 10% FBS at 37°C for 36 h to establish monolayers. Subsequently, the cells were exposed to 200 μL of antibiotic-free DMEM containing PMT at a final concentration of 25 µg/mL and 70 kDa fluorescein isothiocyanate (FITC; Sigma, St. Louis, MO, USA) at a final concentration of 10 µM.

The cells were then incubated for 48 h at 37°C in a 5% CO_2_ environment, following which 100 µL of medium from the basal chamber was transferred to a black-well plate (Greiner Bio-One, Germany). Fluorescence intensity was measured using an excitation/emission wavelength of 490/520 nm in a Victor Nivo multimode plate reader (PerkinElmer, Waltham, MA, USA). Dextran permeability was quantified based on the obtained results, as outlined in previous studies (12).

### Molecular docking and bioinformatical analysis

Molecular docking analysis was conducted to analyze the interactions between PMT and Ca_V_3.1. To achieve this, the three-dimensional structure of Ca_V_3.1 was obtained from the Protein Data Bank (PDB ID: O43497) (22), and predicted three-dimensional structure of PMT was generated using AlphaFold 3, following the published protocols (21). Molecular docking was performed using the protein-protein docking program, PIPER, developed by Schrödinger (New York, US). The ligand was defined with 70,000 rotatable probes, and a total of 30 conformations were generated. PMT was considered as the ligand, while Ca_V_3.1 was designated as the receptor during the docking process. The transmembrane and intracellular segments of the Ca_V_3.1 protein were set as repulsion, while the extracellular segment was set as attraction. A bonus value of 0.99 was applied for constrained docking. Among all the generated conformations, Piper performed clustering of the first 1000 rotational conformations based on the Root Mean Square Deviation (RMSD) between each pair of atoms. The representative conformation for each cluster was selected based on the highest number of neighbors within that cluster. Piper sorted the generated conformations according to the number of clusters in each class, with the top-ranked conformation having the highest cluster count. The first conformation was selected for further analysis. The binding energies were calculated as previously described (36).

### Co-immunoprecipitation (co-IP) assay

To verify the identified binding regions in PMT, a co-immunoprecipitation (co-IP) assay was performed. Initially, a commercially synthesized fragment from nucleotides 2209 to 4506 at the N-terminal of porcine *cacna1g* (GenBank accession no. XM_021067186) was obtained (Tsingke Bio., Beijing, China). This fragment was then cloned into the pcAGGS-N-flag vector to generate the plasmid pcAGGS-N-flag-pCav3.1N_2209-4506_. Subsequently, HEK293T cells were transfected with either pcAGGS-N-flag-pCav3.1N_2209-4506_ or the control vector pcAGGS-N-flag using Lipofectamine 2000 (ThermoFisher, Waltham, US). Following a 48-hour incubation at 37 °C, the cells were washed twice with cold PBS and lysed using a commercial cell lysis buffer for western blot and IP (Beyotime, Shanghai, China) supplemented with protease inhibitor (1:100; Beyotime, Shanghai, China) on ice for 25 minutes. The cell lysates were collected and further subjected to ultrasonication. The resulting mixture was then centrifuged at 4 °C, 12,000 rpm for 10 minutes to collect the supernatants, which were subsequently incubated with DYKDDDDK tag Monoclonal antibody (Proteintech, Wuhan, China) binding rProtein A/G Magarose Beads (Smart-Lifesciences, Changzhou, China) overnight at 4 °C. The beads were washed three times with the cell lysis buffer for western blot and IP and then co-incubated with 3×Flag Peptide (150 μg/mL; Beyotime, Shanghai, China) at 4 °C for 2 hours. Following this, the samples were mixed with 5× protein loading buffer and incubated at 95 °C for 10 minutes. The proteins were separated by SDS-PAGE, transferred onto a polyvinylidene fluoride membrane (Bio-Rad, Hercules, US), and blocked in PBST containing 5% BSA at room temperature for 3 hours. The membrane was then incubated overnight at 4 °C with DYKDDDDK tag Monoclonal antibody (1:5,000; Proteintech, Wuhan, China), His-Tag Monoclonal antibody (1:5,000; Proteintech, Wuhan, China), and/or GAPDH Monoclonal antibody (1:5,000; Proteintech, Wuhan, China). After washing with PBST, the membrane was co-incubated with HRP-conjugated Affinipure Goat Anti-Mouse IgG (H+L) (1:5,000; Proteintech, Wuhan, China) at 37 °C for 1 hour. Finally, the membrane was subjected to enhanced chemiluminescence (ECL) detection using an ECL reagent (Beyotime, Shanghai, China).

### CRISPR/Cas9-based Genome Editing

To knock out the Ca_v_3.1 coding gene (*cacna1g*), three pairs of sgRNAs (cacna1g sg1/sg2/sg3) were designed using CHOPCHOP (https://chopchop.cbu.uib.no/) and synthesized by a commercial company (Tsingke Bio., Beijing, China). The sequences of the sgRNAs can be found in Table S1. The synthesized sgRNAs were cloned into the LentiCRISPR-V2 vector (gifted by Prof. Wentao Li at Huazhong Agricultural University) to generate the plasmids LentiCRISPR-V2-cacna1g sg1/sg2/sg3. Subsequently, the LentiCRISPR-V2-cacna1g sg1/sg2/sg3 plasmids (20 nM) were transfected into NPTr cells using Lipofectamine 2000 (ThermoFisher, Waltham, US). The cells were cultured for an additional 48 hours, followed by the addition of puromycin (3 μg/mL; Beyotime, Shanghai, China) to select for cells with puromycin resistance. Genomic DNAs were extracted from puromycin-resistant cells and used as templates for verifying the knockout of *cacna1g* using the target-specific primers: target1-F/R, target2-F/R, or target3-F/R (Table S1). Knocking out of Ca_v_3.1 was further confirmed using western-blotting. Finally, cells with confirmed knockout profiles were further purified to generate monoclonal cell lines with the *cacna1g* gene knocked out.

### Cytotoxicity tests

The susceptibility of NPTr and MLE-12 cells to PMT was assessed using an MTT Cell Proliferation and Cytotoxicity Assay Kit (Beyotime, Shanghai, China) following the manufacturer’s instructions. Initially, 5000 NPTr or MLE-12 cells per well in a 96-well plate (Corning, Corning, US) were incubated with PMT (final concentration: 25 µg/mL) and the MTT solution (10 μl) for 4 hours at 37 °C under a 5% CO_2_ atmosphere. Subsequently, 100 µL of Formazan solubilization solution was added to each well, followed by continued incubation at 37 °C under a 5% CO_2_ atmosphere for 4 hours. The absorbance at 570 nm was then measured using a BMG Labtech FLUOstar Omega plate reader (BMG Labtech, Allmendgrün, Germany).

Additionally, we evaluated the susceptibility of NPTr wild-type cells and Cav3.1-KO cells to PMT following the CCK-8 protocol (37). In brief, NPTr wild-type cells and Ca_v_3.1-KO cells were seeded into the wells of a 6-well plate (Corning, Corning, US). Cells treated with PBS and Triton X-100 were used as negative and positive controls, respectively. The plate was then incubated at 37 °C under a 5% CO_2_ atmosphere until the cell density reached 90% confluence. Subsequently, the cells were incubated with PMT (100 μg/mL) at 37 °C under a 5% CO_2_ atmosphere for 48 hours. Images were captured using an EVOS M7000 Automatic Living Cell Imaging System (ThermoFisher, Waltham, US). Cell death rates were determined using an Enhanced Cell Counting Kit-8 (CCK-8) assay kit (Beyotime, Shanghai, China) according to the manufacturer’s instructions. The absorbance at OD_450_ was measured using a BMG Labtech FLUOstar Omega plate reader (BMG Labtech, Allmendgrün, Germany). The experiments were independently repeated three times, and the data were subjected to statistical analysis. The cell death rate (%) was calculated using the following formula: Cell death rate (%) = (A_buf_ - A_tox_) / (A_buf_ - A_trtn_) × 100%, where Abuf, Atox, and Atrtn represent the absorbance values of the wells treated with PBS, toxin, and Triton X-100, respectively (37).

### Quantitative real-time PCR (qPCR)

To investigate the influence of PMT on tight junctions and adherens junctions, total RNAs were extracted from the lung tissues of mice inoculated with PMT/PBS (as mentioned in mouse experiments) and/or respiratory epithelial cells treated with PMT/PBS (as mentioned in dextran-based *trans*-well permeability assays). The RNA extraction was performed using the TRIzol reagent (Thermo Fisher, Waltham, MA, USA). Subsequently, the extracted RNAs were used to synthesize cDNA using the PrimeScript RT reagent Kit with gDNA Eraser (TAKARA, Beijing, China). The synthesized cDNA served as the template for determining the transcriptional levels of selected tight junctions (ZO-1, occludin) and adherens junctions (β-catenin, E-cadherin) using the primers listed in Table S1, with GAPDH utilized as the reference gene. The experiments were independently repeated three times.

To understand the effect of PMT on endoplasmic reticulum stress, monolayers of NPTr cells were incubated with 25 µg/mL of PMT for 6 h, 12 h, 24 h, and 48 h, respectively. In additional, cells were also exposed to 5 μM of Ca^+^ chelating agent, BAPTA-AM (MCE, Monmouth Junction, US), for 12 h to investigate the impact of Ca^2+^ on endoplasmic reticulum stress. PBS-treated cells were included as controls. At different time points, both PMT and PBS treated cells were harvested for RNA extraction. The transcription levels of markers (GRP78, CHOP, ATF6, PERK, IRE1) indicating endoplasmic reticulum stress were examined using qPCR with cDNA synthesized from the extracted RNAs as templates, along with the primers listed in Table S1.

### Western blot analysis

Tissue samples (0.1-0.2 g) or cell samples were lysed using RIPA containing protease inhibitor (Beyotime, Shanghai, China). Proteins were quantified using an Enhanced BCA Protein Assay Kit (Beyotime, Shanghai, China). To explore the impact of RhoA/ROCK pathway on the disruption of respiratory barrier induced by PMT, monolayers of NPTr cells were also treated using the ROCK inhibitor Y-27693 (MCE, Monmouth Junction, US) at 10 μm for 48 h. Cells were also treated using 0.1 mM of the endoplasmic reticulum stress inhibitor 4-PBA (MCE, Monmouth Junction, US) for 48 h to investigate the influence of inhibiting endoplasmic reticulum stress on RhoA/ROCK pathway induced by PMT.

Following this, proteins extracted from either murine lung tissues or mammalian respiratory epithelial cells were separated on a 10% SDS-PAGE gel and then transferred onto a PVDF membrane (Bio-Rad, USA). The PVDF membrane was blocked with TBST with 5% BSA for 3 h at room temperature. Subsequently, the membrane was incubated with the following antibodies at the specific dilutions: RhoA monoclonal antibody (1:2000, catalog no. 66733-1-Ig; Proteintech, Wuhan, China), ZO-1 polyclonal antibody (1:5000, catalog no. 21773-1-AP; Proteintech, Wuhan, China), Occludin polyclonal antibody (1:5000, catalog no. 113409-1-AP; Proteintech, Wuhan, China), GAPDH monoclonal antibody (1:20000, catalog no. 60004-1-lg; Proteintech, Wuhan, China), IRE1 polyclonal antibody (1:2000, catalog no. WL02562; Wanleibio, Shenyang, China), p-IRE1 polyclonal antibody (1:2000, catalog no. AF7150; Affinity Biosciences, Liyang, China), GRP78 polyclonal antibody (1:5000, catalog no. 11587-1-AP; Proteintech, Wuhan, China), ROCK1 polyclonal antibody (1:5000, catalog no. 21850-1-AP; Proteintech, Wuhan, China). After washing, the PVDF membrane was incubated with HRP-conjugated Goat Anti-Mouse IgG(H+L) (1:5000, catalog no. SA00001-1; Proteintech, Wuhan, China) or HRP-conjugated Goat Anti-Rabbit IgG(H+L) (1:5000, catalog no. SA00001-2; Proteintech, Wuhan, China), and visualized using the enhanced chemiluminescence (ECL) reagent (Beyotime, Shanghai, China). Protein bands were quantified using ImageJ software.

### Confocal laser scanning microscopy examination

To investigate the influence of PMT treatment on the expression of ZO-1 and changes in the F-actin cytoskeleton in respiratory epithelial cells, NPTr monolayers were inoculated with 25 μ/ml of PMT or PBS for 48 h. Additionally, cells were treated with the ROCK inhibitor Y-27693 (MCE, Monmouth Junction, US) at 10 μm for 48 h to observe the inhibition of RhoA/ROCK signaling on the expression of ZO-1 and changes in the F-actin cytoskeleton induced by PMT. Afterwards, the cells were washed three times with pre-chilled PBS to remove free proteins. Subsequently, the cells were fixed with pre-chilled 4% paraformaldehyde for 2 h and then blocked in 5% BSA at room temperature for 2 h. Following this, the cells were incubated overnight at 4°C with ZO-1 antibody (1:500, catalog no. 21773-1-AP; Proteintech, Wuhan, China). After washing, the cells were incubated in the dark at room temperature with Alexa Fluor™ 488 phalloidin (to stain F-actin; 1:400, catalog no. A12379; Life Technologies, Carlsbad, USA) for 1 h, followed by incubation with goat anti-rabbit IgG (1:500, catalog no. A-21428; Invitrogen, Waltham, USA) in the dark at room temperature for 1 h. Finally, the cells were incubated in the dark at room temperature for 30 minutes with an anti-fade mounting medium containing DAPI (Beyotime, Shanghai, China), and the expression of ZO-1 and changes in the cell cytoskeleton were observed under a superresolution confocal laser scanning microscopy system. The obtained photos were analyzed using NIS-Elements Viewer 4.20 software (Nikon, Tokyo, Japan).

To observe changes in cytoplasmic Ca^2+^ concentration induced by PMT, monolayers of NPTr wild type cells/Ca_V_3.1-KO cells were treated with 25 μg/mL of PMT or PBS for 6 h, 12h, and 24 h, respectively. Afterwards, the cells were washed three times with Hank’s Balanced Salt Solution (HBSS, 8 mg/mL of NaCl, 0.4 mg/mL of KCl, 1 mg/mL of glucose, 60 mg/L of KH_2_PO4, 47.5 mg/L of Na_2_HPO4; pH 7.2) (21). Cells were then incubated with 4 µM of Fluo-3 AM probe (Beyotime, Shanghai, China) for 45 min at 37°C. After washing three times with HBSS, the cells were maintained in the solution for 30 minutes at 37°C. Changes in cytoplasmic Ca2+ concentration were observed under a superresolution confocal laser scanning microscopy system, with an excitation wavelength of 494 nm and an emission wavelength of 516 nm. The obtained photos were analyzed using NIS-Elements Viewer 4.20 software (Nikon, Tokyo, Japan).

### Statistical analysis

Statistical significance for data analysis was determined using Student’s *t-test* in GraphPad Prism 9.0. The data is presented as mean ± SD. The significance level was set at *P* < 0.05 (∗), *P* < 0.01 (∗∗), or *P* < 0.001 (∗∗∗). The term “ns” indicates no significant difference.

## Supplemental materials

**Table S1.** Primers and sequences of sgRNAs used in this study.

**Fig.S1.** Generation of NPTr *cacna1g* knockout cells. Panel A shows the construction of *cacna1g* knockout NPTr cells by using the CRISPR/Cas9-based genome editing. Panels B and C displays the susceptibility of NPTr wild type cells and *cacna1g* knockout cells to PMT. Data represent mean ± SD. The significance level was set at *P* < 0.001 (∗∗∗).

## Acknowledgments

We acknowledge Dr. Wentao Li at Huazhong Agricultural University for the gift of LentiCRISPR-V2, and Dr. Mengjia Zhang for the help on CRISPR-Cas9 editing technology. This study was funded in part by Hubei Provincial Natural Foundation of China (No. 2023AFA094), Yingzi Tech & Huazhong Agricultural University Intelligent Research Institute of Food Health (No. IRIFH202209), the Fundamental Research Funds for the Central Universities (No. 2662023PY005), Walmart Foundation (Project # 61626817), and Hubei Hongshan Laboratory & Huazhong Agricultural University Startup fund. The funder had no role in the study design, data collection, data analysis, data interpretation, or writing of the manuscript.

## Reference

1. Wilson BA, Ho M. 2013. Pasteurella multocida: from zoonosis to cellular microbiology. Clin Microbiol Rev 26:631–55.

2. WOAH. 2018. Atrophic rhinitis of swine. OIE Terrestrial Manual 2018 Chapter 3.8.2.:1540–1550.

3. Riising HJ, van Empel P, Witvliet M. 2002. Protection of piglets against atrophic rhinitis by vaccinating the sow with a vaccine against Pasteurella multocida and Bordetella bronchiseptica. Vet Rec 150:569–71.

4. Kloos B, Chakraborty S, Lindner SG, Noack K, Harre U, Schett G, Krämer OH, Kubatzky KF. 2015. Pasteurella multocida toxin-induced osteoclastogenesis requires mTOR activation. Cell Commun Signal 13:40.

5. Kitadokoro K, Kamitani S, Miyazawa M, Hanajima-Ozawa M, Fukui A, Miyake M, Horiguchi Y. 2007. Crystal structures reveal a thiol protease-like catalytic triad in the C-terminal region of Pasteurella multocida toxin. Proc Natl Acad Sci U S A 104:5139–44.

6. Orth JH, Aktories K. 2012. Molecular biology of Pasteurella multocida toxin. Curr Top Microbiol Immunol 361:73–92.

7. Wilson BA, Ho M. 2012. Pasteurella multocida toxin interaction with host cells: entry and cellular effects. Curr Top Microbiol Immunol 361:93–111.

8. Schoellkopf J, Mueller T, Hippchen L, Mueller T, Reuten R, Backofen R, Orth J, Schmidt G. 2022. Genome wide CRISPR screen for Pasteurella multocida toxin (PMT) binding proteins reveals LDL Receptor Related Protein 1 (LRP1) as crucial cellular receptor. PLoS Pathog 18:e1010781.

9. Buckley A, Turner JR. 2018. Cell Biology of Tight Junction Barrier Regulation and Mucosal Disease. Cold Spring Harb Perspect Biol 10:a029314.

10. Yuksel H, Turkeli A. 2017. Airway epithelial barrier dysfunction in the pathogenesis and prognosis of respiratory tract diseases in childhood and adulthood. Tissue Barriers 5:e1367458.

11. Lin L, Yang J, Zhang D, Lv Q, Wang F, Liu P, Wang M, Shi C, Huang X, Liang W, Tan C, Wang X, Chen H, Wilson BA, Wu B, Peng Z. 2023. Vascular Endothelial Growth Factor A Contributes to Increased Mammalian Respiratory Epithelial Permeability Induced by Pasteurella multocida Infection. Microbiol Spectr 11:e0455422.

12. Zhang D, Lin L, Yang J, Lv Q, Wang M, Hua L, Zhang K, Chen H, Wu B, Peng Z. 2024. Pseudorabies virus infection increases the permeability of the mammalian respiratory barrier to facilitate Pasteurella multocida infection. mSphere 9:e0029724.

13. Li L, Xin J, Wang H, Wang Y, Peng W, Sun N, Huang H, Zhou Y, Liu X, Lin Y, Fang J, Jing B, Pan K, Zeng Y, Zeng D, Qin X, Bai Y, Ni X. 2023. Fluoride disrupts intestinal epithelial tight junction integrity through intracellular calcium-mediated RhoA/ROCK signaling and myosin light chain kinase. Ecotoxicol Environ Saf 257:114940.

14. Li B, Zhao WD, Tan ZM, Fang WG, Zhu L, Chen YH. 2006. Involvement of Rho/ROCK signalling in small cell lung cancer migration through human brain microvascular endothelial cells. FEBS Lett 580:4252–60.

15. Mueller BK, Mack H, Teusch N. 2005. Rho kinase, a promising drug target for neurological disorders. Nat Rev Drug Discov 4:387–98.

16. Luo S, Ho M, Wilson BA. 2008. Application of intact cell-based NFAT-beta-lactamase reporter assay for Pasteurella multocida toxin-mediated activation of calcium signaling pathway. Toxicon 51:597–605.

17. Kubatzky KF. 2022. Pasteurella multocida toxin - lessons learned from a mitogenic toxin. Front Immunol 13:1058905.

18. Teruya S, Hiramatsu Y, Nakamura K, Fukui-Miyazaki A, Tsukamoto K, Shinoda N, Motooka D, Nakamura S, Ishigaki K, Shinzawa N, Nishida T, Sugihara F, Maeda Y, Horiguchi Y. 2020. Bordetella Dermonecrotic Toxin Is a Neurotropic Virulence Factor That Uses Ca(V)3.1 as the Cell Surface Receptor. mBio 11:e03146–19.

19. Lizák B, Birk J, Zana M, Kosztyi G, Kratschmar DV, Odermatt A, Zimmermann R, Geiszt M, Appenzeller-Herzog C, Bánhegyi G. 2020. Ca(2+) mobilization-dependent reduction of the endoplasmic reticulum lumen is due to influx of cytosolic glutathione. BMC Biol 18:19.

20. Daverkausen-Fischer L, Pröls F. 2022. Regulation of calcium homeostasis and flux between the endoplasmic reticulum and the cytosol. J Biol Chem 298:102061.

21. Abramson J, Adler J, Dunger J, Evans R, Green T, Pritzel A, Ronneberger O, Willmore L, Ballard AJ, Bambrick J, Bodenstein SW, Evans DA, Hung CC, O’Neill M, Reiman D, Tunyasuvunakool K, Wu Z, Žemgulytė A, Arvaniti E, Beattie C, Bertolli O, Bridgland A, Cherepanov A, Congreve M, Cowen-Rivers AI, Cowie A, Figurnov M, Fuchs FB, Gladman H, Jain R, Khan YA, Low CMR, Perlin K, Potapenko A, Savy P, Singh S, Stecula A, Thillaisundaram A, Tong C, Yakneen S, Zhong ED, Zielinski M, Žídek A, Bapst V, Kohli P, Jaderberg M, Hassabis D, Jumper JM. 2024. Accurate structure prediction of biomolecular interactions with AlphaFold 3. Nature 630:493–500.

22. Zhao Y, Huang G, Wu Q, Wu K, Li R, Lei J, Pan X, Yan N. 2019. Cryo-EM structures of apo and antagonist-bound human Ca(v)3.1. Nature 576:492–497.

23. Zihni C, Mills C, Matter K, Balda MS. 2016. Tight junctions: from simple barriers to multifunctional molecular gates. Nat Rev Mol Cell Biol 17:564–80.

24. Harris TJ, Tepass U. 2010. Adherens junctions: from molecules to morphogenesis. Nat Rev Mol Cell Biol 11:502–14.

25. Dudet LI, Chailler P, Dubreuil JD, Martineau-Doize B. 1996. Pasteurella multocida toxin stimulates mitogenesis and cytoskeleton reorganization in Swiss 3T3 fibroblasts. J Cell Physiol 168:173–82.

26. Essler M, Hermann K, Amano M, Kaibuchi K, Heesemann J, Weber PC, Aepfelbacher M. 1998. Pasteurella multocida toxin increases endothelial permeability via Rho kinase and myosin light chain phosphatase. J Immunol 161:5640–6.

27. Blum MS, Toninelli E, Anderson JM, Balda MS, Zhou J, O’Donnell L, Pardi R, Bender JR. 1997. Cytoskeletal rearrangement mediates human microvascular endothelial tight junction modulation by cytokines. Am J Physiol 273:H286–94.

28. Fanning AS, Jameson BJ, Jesaitis LA, Anderson JM. 1998. The tight junction protein ZO-1 establishes a link between the transmembrane protein occludin and the actin cytoskeleton. J Biol Chem 273:29745–53.

29. Feng S, Zou L, Wang H, He R, Liu K, Zhu H. 2018. RhoA/ROCK-2 Pathway Inhibition and Tight Junction Protein Upregulation by Catalpol Suppresses Lipopolysaccaride-Induced Disruption of Blood-Brain Barrier Permeability. Molecules 23:2371.

30. He F, Peng J, Deng XL, Yang LF, Wu LW, Zhang CL, Yin F. 2011. RhoA and NF-κB are involved in lipopolysaccharide-induced brain microvascular cell line hyperpermeability. Neuroscience 188:35–47.

31. Dopeso H, Rodrigues P, Cartón-García F, Macaya I, Bilic J, Anguita E, Jing L, Brotons B, Vivancos N, Beà L, Sánchez-Martín M, Landolfi S, Hernandez-Losa J, Ramon YCS, Nieto R, Vicario M, Farre R, Schwartz S, Jr., van Ijzendoorn SCD, Kobayashi K, Martinez-Barriocanal Á, Arango D. 2024. RhoA downregulation in the murine intestinal epithelium results in chronic Wnt activation and increased tumorigenesis. iScience 27:109400.

32. Krebs J, Agellon LB, Michalak M. 2015. Ca(2+) homeostasis and endoplasmic reticulum (ER) stress: An integrated view of calcium signaling. Biochem Biophys Res Commun 460:114–21.

33. Haws HJ, McNeil MA, Hansen MD. 2016. Control of cell mechanics by RhoA and calcium fluxes during epithelial scattering. Tissue Barriers 4:e1187326.

34. Liu W, Yang M, Xu Z, Zheng H, Liang W, Zhou R, Wu B, Chen H. 2012. Complete genome sequence of Pasteurella multocida HN06, a toxigenic strain of serogroup D. J Bacteriol 194:3292–3.

35. Percie du Sert N, Hurst V, Ahluwalia A, Alam S, Avey MT, Baker M, Browne WJ, Clark A, Cuthill IC, Dirnagl U, Emerson M, Garner P, Holgate ST, Howells DW, Karp NA, Lazic SE, Lidster K, MacCallum CJ, Macleod M, Pearl EJ, Petersen OH, Rawle F, Reynolds P, Rooney K, Sena ES, Silberberg SD, Steckler T, Würbel H. 2020. The ARRIVE guidelines 2.0: Updated guidelines for reporting animal research. PLoS Biol 18:e3000410.

36. Kukol A. 2008. Molecular Modeling of Proteins. Meth Mol Bio 443:365–382.

37. Yuan J, Zhao Q, Li J, Wen Y, Wu R, Zhao S, Lang YF, Yan QG, Huang X, Du S, Cao SJ. 2024. CXCL8 Knockout: A Key to Resisting Pasteurella multocida Toxin-Induced Cytotoxicity. Int J Mol Sci 25:5330.

